# Computational structure prediction methods enable the systematic identification of oncogenic mutations

**DOI:** 10.1101/2022.11.18.517015

**Authors:** Xi Fu, Clara Reglero, Vinay Swamy, Jui Wan Loh, Hossein Khiabanian, Robert Albero, Farhad Forouhar, Mohammed AlQuraishi, Adolfo A. Ferrando, Raul Rabadan

**Affiliations:** Program for Mathematical Genomics, Department of Systems Biology, Columbia University, New York, NY, USA; Department of Biomedical Informatics, Columbia University, New York, NY, USA; Institute for Cancer Genetics, Columbia University, New York, NY, USA; Rutgers Cancer Institute of New Jersey, Rutgers University, New Brunswick, NJ, 08901, USA; Center for Systems and Computational Biology, Rutgers Cancer Institute of New Jersey, Rutgers University, New Brunswick, NJ, 08901, USA; Department of Pathology and Laboratory Medicine, Rutgers Robert Wood Johnson Medical School, Rutgers University, New Brunswick, NJ, 08901, USA; Proteomics and Macromolecular Crystallography Shared Resource, Herbert Irving Comprehensive Cancer Center, Columbia University, New York, NY, USA; Department of Pediatrics, Columbia University, New York, NY, USA

## Abstract

Oncogenic mutations are associated with the activation of key pathways necessary for the initiation, progression and treatment-evasion of tumors. While large genomic studies provide the opportunity of identifying these mutations, the vast majority of variants have unclear functional roles presenting a challenge for the use of genomic studies in the clinical/therapeutic setting. Recent developments in predicting protein structures enable the systematic large-scale characterization of structures providing a link from genomic data to functional impact. Here, we observed that most oncogenic mutations tend to occur in protein regions that undergo conformation changes in the presence of the activating mutation or when interacting with a protein partner. By combining evolutionary information and protein structure prediction, we introduce the Evolutionary and Structure (ES) score, a computational approach that enables the systematic identification of hotspot somatic mutations in cancer. The predicted sites tend to occur in Short Linear Motifs and protein-protein interfaces. We test the use of ES-scores in genomic studies in pediatric leukemias that easily recapitulates the main mechanisms of resistance to targeted and chemotherapy drugs. To experimentally test the functional role of the predictions, we performed saturated mutagenesis in NT5C2, a protein commonly mutated in relapsed pediatric lymphocytic leukemias. The approach was able to capture both commonly mutated sites and identify previously uncharacterized functionally relevant regions that are not frequently mutated in these cancers. This work shows that the characterization of protein structures provides a link between large genomic studies, with mostly variants of unknown significance, to functional systematic characterization, prioritizing variants of interest in the therapeutic setting and informing on their possible mechanisms of action.

## Introduction

Mutations in proteins have a major role in the onset and development of cancer and treatment resistance. The special role of mutations is determined by the diversity of their impact on molecular function. Large-scale genomic approaches are unveiling a large number of mutations, the vast majority of which have unclear functional consequences. In this context, the systematic identification of functional oncogenic mutations in cancer has led to the development of computational methods that include recurrence-based^1,2^ and model-based approaches^3^. Recurrence-based approaches compare the observed mutation rate with a background model on a single residue^1,2^, a fixed window of sequence^4,5^, or a cluster in 3D structure^6,7^. Model-based approaches^8,9^, largely benchmarked in recent studies^10,11^, use biochemical, evolutionary, or structural features to build predictive models or scores to distinguish oncogenic mutations from passenger ones. However, both methods use information from recurrent cancer mutations, which could potentially bias the mechanistic interpretation of the model. Unsupervised methods^12–14^ that do not directly rely on recurrence information might provide better biological insights on cancer hotspots and a more unbiased way to identify less common cancer risk variants.

Protein structure provides a link between genetic changes and functional implications. Unlike inactivating mutations that may occur across the genome, driver mutations localize at specific sites within the oncogenic proteins and are associated with the gain of specific functions leading to the activation of key cancer-related pathways^1^. For instance, mutations in the v-raf murine sarcoma viral oncogenes homolog B1 (BRAF) protein, commonly found across many cancers including melanomas, brain tumors, and some leukemias, activate the MAP kinase/ERKs signaling pathway, a key pathway involved in a variety of cancer processes including proliferation and cell cycle progression. Most BRAF oncogenic mutations occur in the V600 amino acid leading to constitutive activation of the kinase and a lack of response to negative feedback mechanisms^15^. Like with BRAF, many oncogenic mutations tend to be located in specific sites including phosphorylation sites^16^, protein-protein interfaces^17^, or intrinsically disordered regions^18^. The incorporation of structure-based methods may not only help with the identification of potential oncogenic mutations but also characterize the potential functional mechanisms.

The recent development of methods to computationally predict protein structures provide a way to systematically incorporate structural information into genomic analyses. Here we introduce the <u>E</u>volutionary and <u>S</u>tructure (ES) score, a simple unsupervised scoring algorithm that combines evolutionary and structural information to identify potential functional sites in human proteins mutated in cancer. We show that ES score outperforms current state-of-the-art computational methods at identifying mutation hotspots in oncogenes. Applying ES score to study longitudinal acute lymphoblastic lymphoma (ALL) genomic studies, we successfully capture the main mechanisms of resistance to 6-mercaptopurine (6-MP) chemotherapy and identify a novel regulatory region of NT5C2 nucleotidase activity leading to chemotherapy resistance.

## Results

### The ES score identifies oncogenic mutations affecting functional protein regions

Our ES score takes advantage of both evolutionary and structural information (Fig 1A). Evolutionary conservation has been used to identify functional regions in proteins. Recent advances on representation learning and large pretrained language models allow to embed millions of peptide sequences found in nature into high dimensional spaces that preserve the evolutionary constraint properties. These models can be used to evaluate the likelihood of observing a specific amino acid at a given position in a protein sequence^19^. However, such evolutionary embeddings might not directly reflect structure properties. On the other hand, protein structure models like Alphafold^20^ can predict structures and provide reliability of the prediction as encoded by the per-residue confidence score (pLDDT). Regions predicted with high confidence tend to be in the structurally conserved core of the protein while low confidence regions are mostly associated with flexible loops that undergo conformational changes when physically interacting with other proteins. We hypothesize that gain-of-function mutations may localize in the transition between ordered and disorder structures as these mutations have been shown to lock conformational structures. To quantify the presence of these ordered/disorder regions we 1) look at the derivative of the pLDDT score and 2) multiply the evolutionary embedding-based variant impact score. By multiplying the two components together and smoothing over 3D structures, our ES score is an unsupervised scoring algorithm that is easily interpretable and widely generalizable (See Methods).

**Figure 1.**
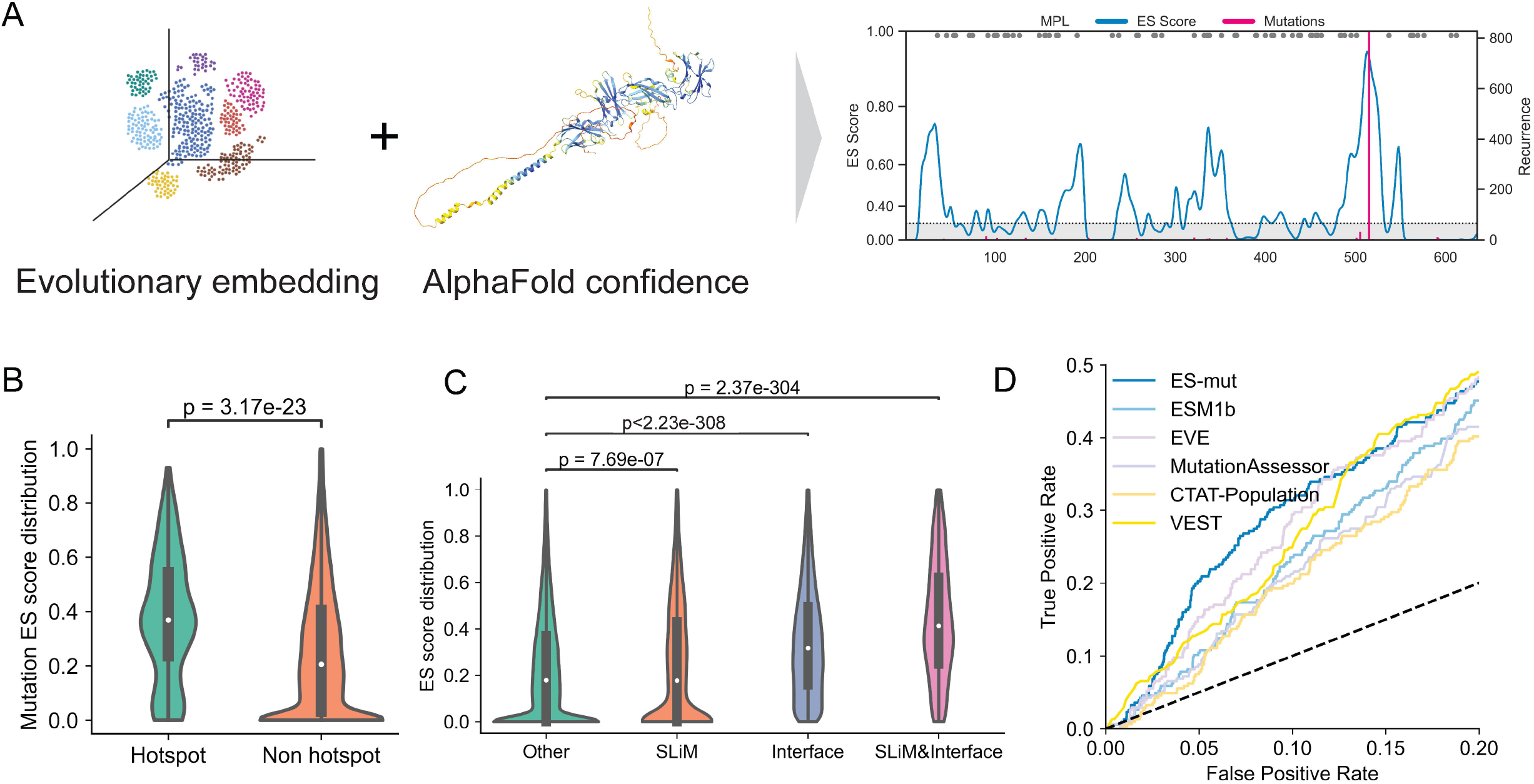
ES score identifies hotspot oncogenic mutations in cancer without supervision. A) ES score is defined using both deep learning-based embedding of evolutionary information and AlphaFold-predicted structure and confidence information. ES score plot for MPL protein sequences is shown on the right as an example. Blue curve marks the smoothed ES score. Red vertical lines marks cancer somatic mutations annotated in COSMIC database. Grey dots are residues prune to mutation based on mutation probability analysis. Light grey shads makes the average ES score across all residues in a protein. B) Violin plot of ES score distribution of cancer hotspot or non hotspot mutations. Welch’s t-test P-value 3.17e-23. Median for hotspots is 0.36, median for non-hotspots is 0.20. C) ES score distribution across oncogene residues classified based on overlap with short linear motif (SliM) and protein-protein interaction interface. Welch’s t-test of SliM v.s. Other: P=7.68e-07; Interface v.s. Other: P<2.23e-308; SliM&Interface v.s. Other: P=2.37e-304. Median of the four groups: Other=0.18, SliMs=0.18, Interface=0.32, SliM&Interface=0.41. D) Receiver operating curve comparison of different methods for oncogenic mutation discovery. The FDR<0.2 region is shown on the plot. Yellow colors marks supervised methods. Blue and purple colors represents unsupervised methods. P-value from Welch’s T test is shown for panel B and C.

### The ES score effectively predicts hotspot mutations affecting protein functionality in cancer

To evaluate whether structural information helps to identify hotspot mutations in oncogenes, we collected all somatic mutations in the COSMIC database^21^ and separated them into two groups: hotspots (HS) which account for more than 10% of all mutation recurrence in a gene, and non-hotspots (NHS), which cover the rest of the mutated residues. Focusing on annotated oncogenes carrying gain-of-function mutations, we found that oncogene mutation hotspots tend to have higher ES scores than non-hotspots (Fig. 1B). Interestingly, we found these higher scores tend to occur in Short Linear Motifs (SLiMs) that span phosphorylation sites and protein-protein interaction interfaces (Fig. 1C). When using a mutation-specific version of ES score that better accounts for amino-acid properties (ES-mut, see Method), the performance increases significantly over both position-specific ES-score and other ESM-based mutation-specific scores (Supp. Fig. 1), suggesting the relevance of both structure and evolutionary information. Altogether, these results support the hypothesis that cancer hotspot mutation tends to hit residues that are involved in conformational regulation through post-translational modification or protein-protein interaction and demonstrate the potential use of ES scores to predict cancer hotspot mutations.

### The ES score outperforms standard supervised and unsupervised methods

We benchmarked ES score against existing methods that predict oncogenic mutations using a list of TCGA somatic mutations^8^ in oncogenes. To avoid information leakage, we selected 4 other methods that are not specifically trained on cancer mutation data: EVE^12^, ESM1b^13,14^, and MutationAssessor^9^ are unsupervised methods while CTAT-Population^8^ and VEST^3^ are supervised methods. For assessing the hotspot mutations in cancer, we are interested in performance with a low false discovery rate as the functional characterization of each potential hotspot will be expensive. In this regime, our ES score achieved the highest performance when controlling the false discovery rate. At a 5% false discovery rate, our method achieves a true positive rate (TPR) of 20.7%, while the second-best method is EVE with a 14.6% TPR (Fig. 1D).

### Mutations with high ES are enriched in genes and sites recurrently mutated in cancer

To further explore the relationship of high ES score with cancer mutation recurrence, we compute the average ES score for the protein sequence of each oncogene across all residues and COSMIC samples. The top 20 oncogenes with the highest average ES scores are shown in Figure 2A. Interestingly, we observe a significant correlation between the score and gene mutation recurrence (Figure 2B), suggesting that the mutations in protein site highlighted by high ES scores are enriched in cancer. The list includes well-known cancer driver proteins, including, RHOA, AKT1, IDH1, IDH2, BRAF, EGFR, FOXL2, RRAC1, PRKACA, NRAS, HRAS, KIT, and KRAS. The per-residue ES scores for the top four proteins are shown in Figure 2C. For example, the mutation hotspot in IDH1 is R132, which is on the boundary between the protein’s small domain and clasp domain and forms an active center with T77, S94, R100, R109, S278, all of which have high ES scores. Another example is IDH2, which host two mutation hotspots, R140 and R172, both have high ES score and forms the active center of the protein in 3D structures. These results demonstrate the potential of using ES score to identify novel functional regions associated with cancer mutations. In addition, the list includes some proteins not well reported in recurrence-based cancer studies. For example, RRAS2, a GTPase that is mutated at low frequencies in ALL, ovarian cancer and melanoma. In particular codon 72 have been functionally validated *in vivo* as a potent oncogenic driver^22^.

**Figure 2.**
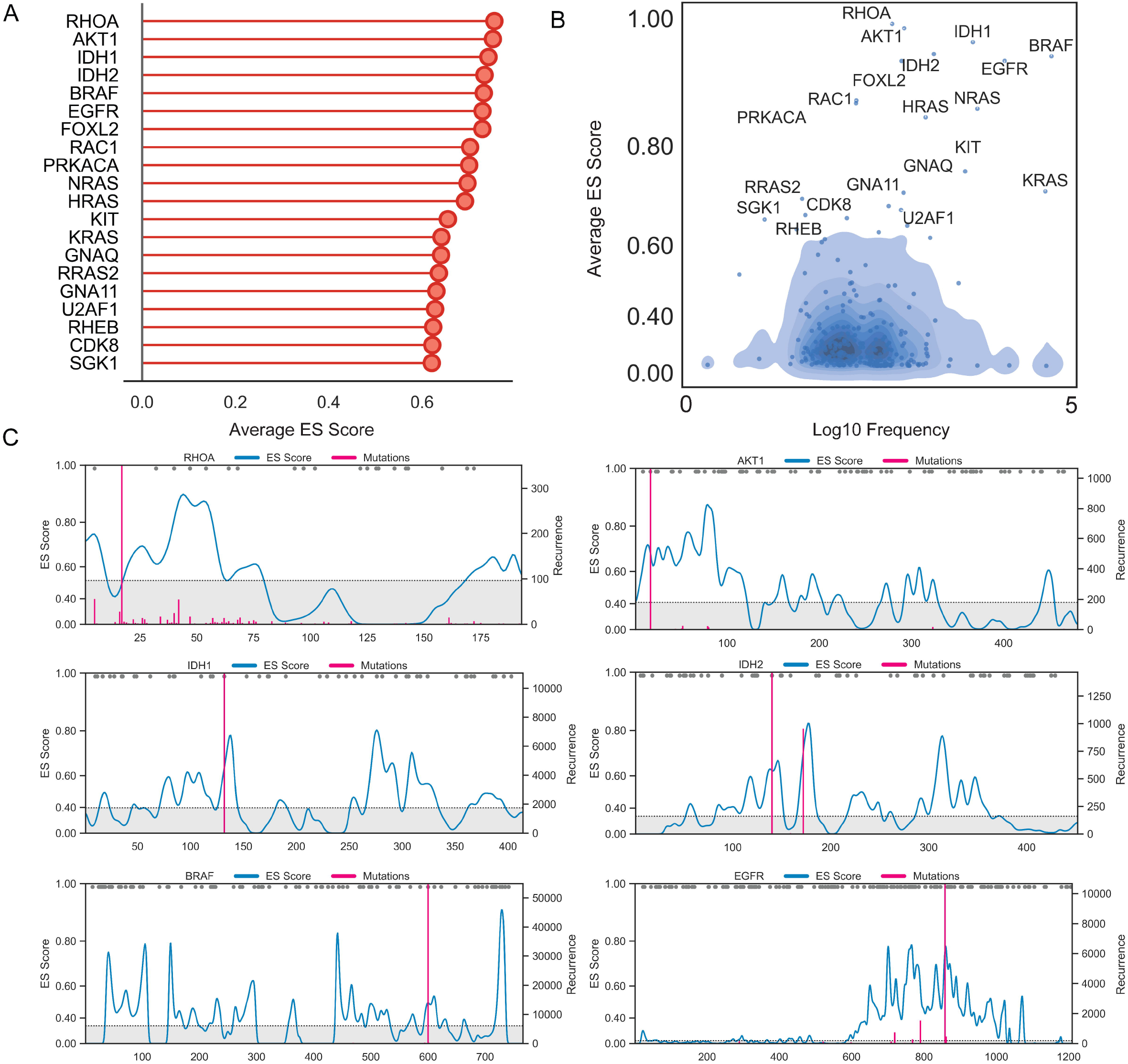
Average ES score at missense mutation sites across 316,701 COSMIC samples. **5)** Top 20 oncogenes ranked by average ES score. B) Average ES scores are associaed with oncogene mutation frequency in COSMIC samples. Blue shades marks the expected distribution using bootstraping (Methods). The Spearman’s correlation between all ‘unexpected’ proteins is 0.46 (P-value=3.14e-05). C) Visualization of ES score for top six proteins in A). Color legends follows that in Figure 1A.

### Common mutations in pediatric and adult acute lymphoblastic leukemias

To test the ability of the ES Score to identify new regions with potential functional roles we focused on ALL, a tumor with known gain-of-function mutations acquired in the course of therapy. With this purpose we collected mutation longitudinal profiles of diagnosed and relapsed ALL samples from a previous study^23^ (Figure 3A) and we computed ES scores for respective proteins with potential oncogenic recurrent mutations both at diagnosis and relapse (Figure 3B). In addition to recapitulating the known frequent drivers of the disease (KRAS, NRAS, BRAF), the ES scores identify relapse-specific mutations that indicate a potential gain-of-function associated with therapeutic resistance in *ABL1* and *NT5C2. BCR-ABL1* fusions occur in 3-5% of pediatric acute lymphoblastic leukemias and are typically associated with worse prognosis^24^. These cases are treated with tyrosine kinase inhibitors (TKI) in addition to conventional chemotherapy^24,25^. Importantly, relapse-specific ABL1 mutations occur in the kinase domain of BCR-ABL1 relapsed leukemias treated with TKIs^23^, corresponding to the region of highest ES scores in the ABL protein (Figure 3C). In particular, the full-length BCR-ABL1 p190 isoform protein structure predicted using AlphaFold2 reveals a close 3D distance between the highest ES score region and the BCR fusion segment. As another example, relapse-specific gain-of-functions mutations in the 5’-nucleotidase, cytosolic II (*NT5C2*) gene induce resistance to 6-MP chemotherapy agent^26,27^. Relapse-associated mutant forms of NT5C2 increase the export of purines to the extracellular matrix and inactivate the cytotoxic metabolites of 6-MP, driving resistance to this drug. NT5C2 functions as a dimer of dimers, with the dimer being the smallest functional unit with enzymatic activity. Therefore, we computed the ES score for the dimeric and monomeric forms (Fig. 3D, left) and we observed that NT5C2 mutations in relapsed leukemias concentrate in regions with high ES score (Fig. 3D) including already characterized domains like the helix A catalytic center (K359, L375), the intermonomeric pocket (R39, R238, R367, among others), and the arm region (K404, D407, S408, P414, and D415)^26,27^; which the dimeric ES score better recapitulates these functional regions. Interestingly, we also detected a region (450-500aa) showing high ES scores with a paucity of mutations. Mutation probability analysis (Method) using COSMIC somatic mutation data shows low probability of mutation compared to the three hotspots (R39, R238, R367, Supp. Fig. 2). This observation indicates that frequency of recurrent mutations is determine by both potential functional selection (ES score) and tumor-associated mutagenesis process (mutation probability).

**Figure 3.**
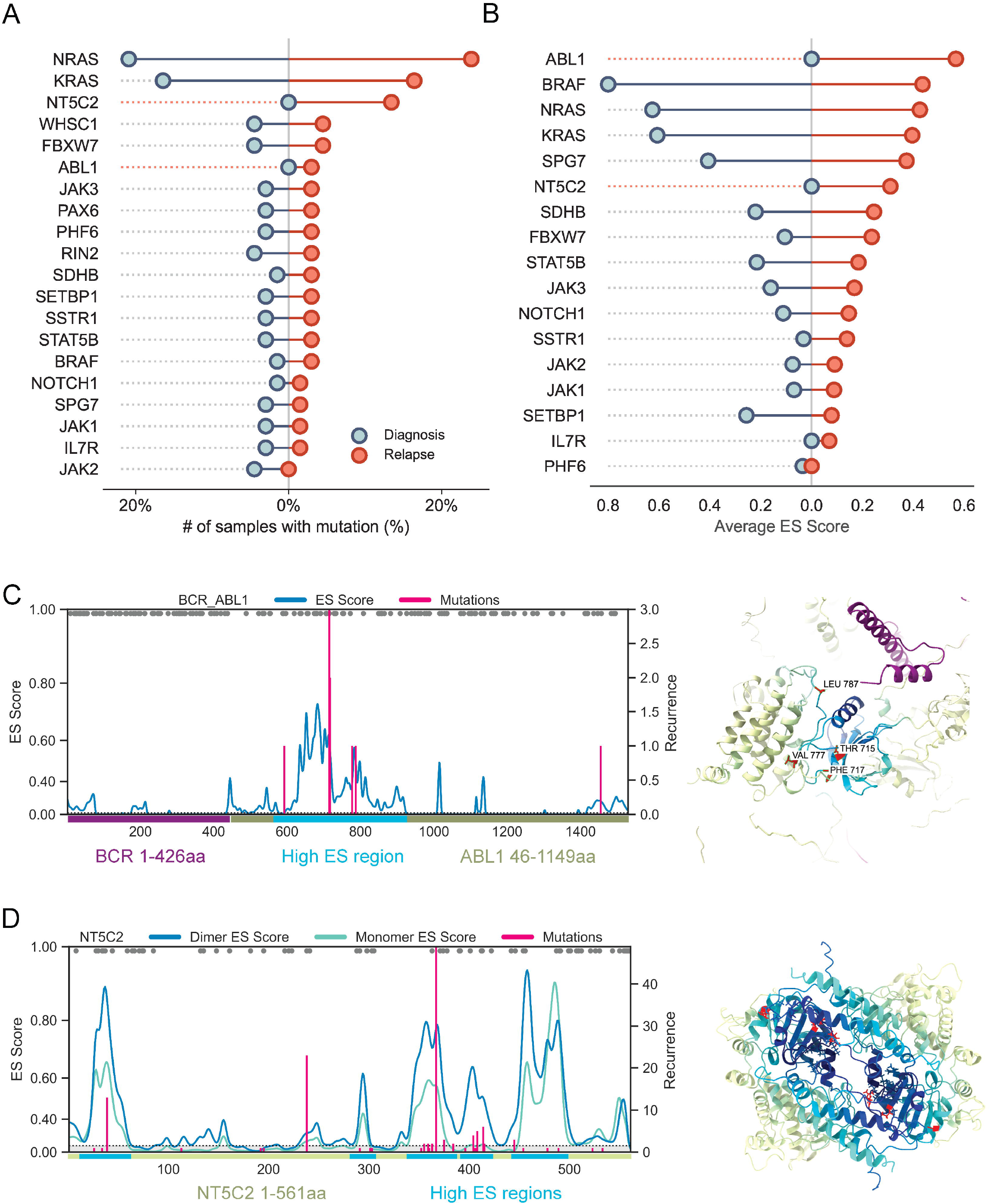
Study of somatic mutations in acute lymphblastic lymphoma with ES score. **5)** Top genes ranked by mutation frequency in acute lymphblastic lymphoma. X axis show mutation frequency in Diagnosis (blue) or Relapse (red) samples. B) Top genes ranked by average ES scores. X axis show average ES scores in Diagnosis (blue) or Relapse (red) samples. C) ES score visualization of *BCR-ABL1* fusion protein and corresponding AlphaFold2 predicted structures. Color legends for ES score follows Figure 1A. Purple region highlights 1-426aa from *BCR*, blue region highlights the high ES score region in the protein, yellow-green region shows the rest of amino acids from *ABL1* 46-1149aa. Mutation hotspots (THR715, PHE717, VAL777, LEU787 are highlighed in red). D). ES score visualization of *NT5C2* monomer (green curve) or dimer (blue curve) and AlphaFold2 predicted *NT5C2* dimer structure. Blue region shows high ES regions in protein seuqnces and structure.

### ES score profile correlates with saturated mutagenesis of NT5C2

In order to systematically identify gain-of-function NT5C2 alleles, map protein domains regulating enzymatic activity and driving resistance to 6-MP and experimentally verify our observations predicted by the ES Score analysis, we generated a saturation mutagenesis library comprising 10,659 variants in which each codon in NT5C2 was systematically mutated to generate all possible amino acid substitutions. Next, we transduced Jurkat ALL cells with a multiplicity of infection of 0.8 and treated this pool with 6-MP at the calculated seven-day IC90 to select for resistant alleles (Supp. Fig 3A). The allelic composition of library-encoded NT5C2 in the resulting resistant (6-MP selected) and control (vehicle only treated) populations were characterized by deep sequencing (Fig. 4A). These analyses revealed a distribution of NT5C2 resistant alleles which are highly consistent with the pattern of ES score (Fig. 3D and Fig. 4B) as well as resistance-driving relapse-associated NT5C2 mutations recovered from genomic profiling studies on relapsed ALL (Fig. 4C). In all, 68% of residues clinically mutated in relapsed ALL were enriched after 6-MP selection in our screen (Supp. Fig 3B). Consistently, we observed a clear enrichment of 6-MP resistant alleles in protein domains critical for NT5C2 nucleotidase activity regulation, including the helix A region and the inhibitory arm domain (Fig. 4D). Moreover, and most interestingly, we observed a high density of NT5C2 mutations in two additional regions: a stretch of nine amino acids (residues 470-478) preceding the C-terminus acidic tail domain and matching with the region pointed by our ES Score, and a segment of seventeen amino acids (residues 26-42) in the N-terminal domain of the NT5C2 protein (Fig. 4E). Structural analysis of these candidate new regulatory regions in the basal and active NT5C2 structures revealed that the 470-478 region forms a hydrophobic pocket located on the surface of the NT5C2 protein (Fig. 5A, B). Interestingly, this region undergoes a dynamic switch-off interactions accommodating the inhibitory arm domain loop of NT5C2 in the basal inactive state and the N-term domain of the protein in the active protein conformation. The potential importance of this mechanism in the regulation of NT5C2 activity is highlighted by the role of the arm domain loop as a switch-off mechanism targeted by recurrent activating mutations in relapse ALL. Based on these results, we hypothesized that the integrity of the 470-478 region may be required for the proper localization and activity of the NT5C2 arm domain loop and that a functional interaction between the N-terminal domain and this region may also participate in promoting the return of NT5C2 to its inactive basal configuration following allosteric activation. To test if these newly identified candidate regulatory regions can influence NT5C2 nucleotidase activity and tumor cell sensitivity to 6-MP, we individually expressed three distinct 470-478 NT5C2 alleles (P472Q, F473P, R478H) and three different N-terminal region mutations (Y27M, R34K, V37W) and analyzed their impact in the sensitivity of Jurkat cells to 6-MP in cellular viability assays (Fig. 5C). Across these analyses, each of these mutations induced increased resistance to 6-MP in a similar extent that the most frequent relapse-associated mutation R367Q compared to wild-type NT5C2 (Fig. 5D and E). These results implicate the 470-478 and N-terminal regions as new regulatory domains of NT5C2 enzymatic activity with potential effects in the response to 6-MP treatment. In addition, these findings further validate the ES Score as a powerful analysis tool for the identification of alleles with oncogenic potential.

**Figure 4.**
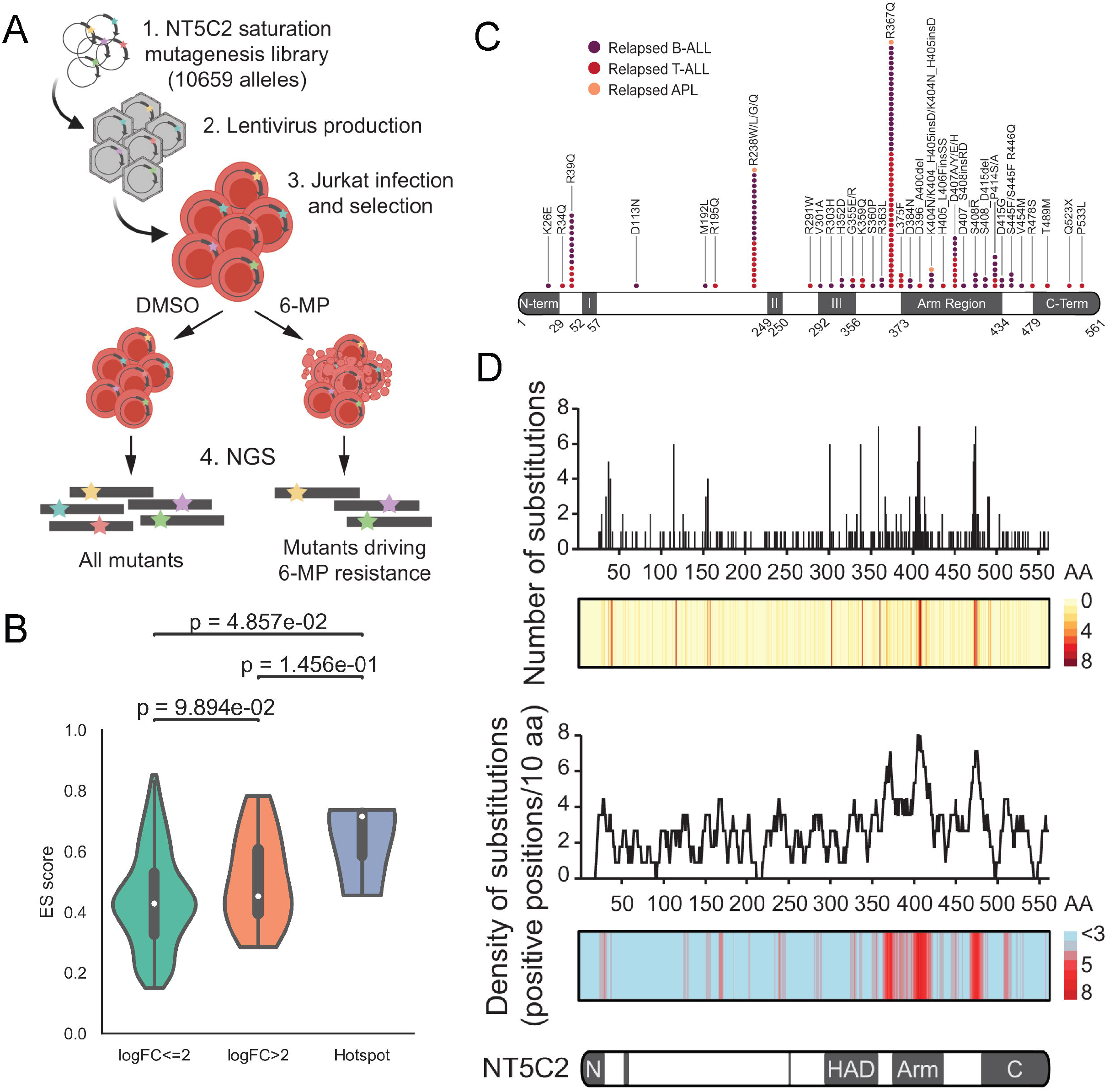
NT5C2 saturation mutagenesis validates ES score profile and identifies new regulatory regions. A) Schematic illustration of NT5C2 saturation mutagenesis screening in Jurkat T-ALL cells. B) ES score distribution across non-hotspot residue with logFC<=2, logFC>2 and hotspot residues. Welch’s T test was performed to compare between all groups. P values are shown above the plot. C) Graphical representation of relapse-associated mutations in B-ALL, T-ALL and APL. HAD core motifs are shown in gray rectangles with roman numerals. D) Number and density of NT5C2 substitutions conferring resistance to 6-MP. Density of substitutions represents the average number of positions where we find significantly resistant mutations (logFDR<4), calculated within a sliding window of 10 amino acids.

**Figure 5.**
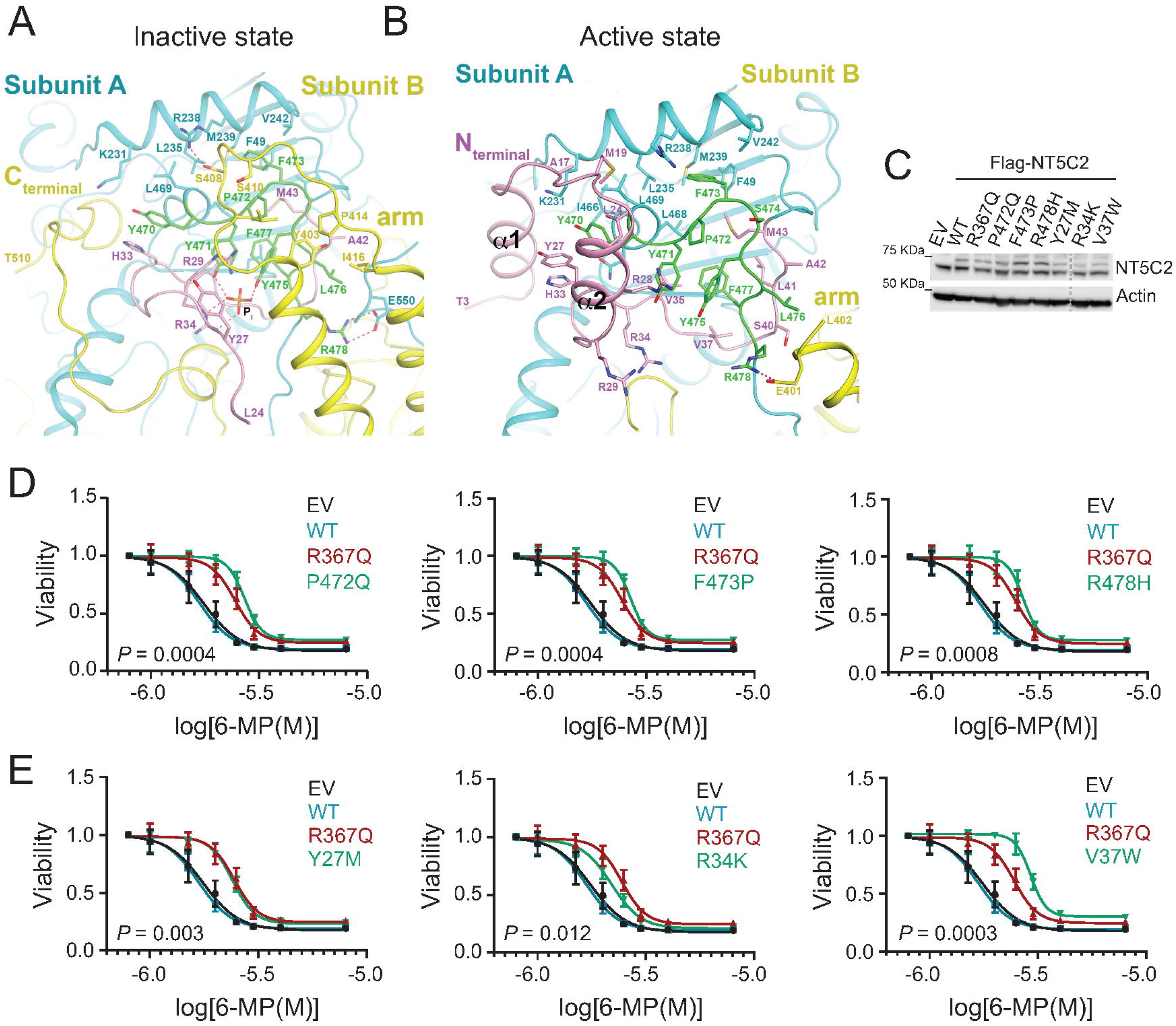
470-478 hydrophobic pocket and the N-terminal constitute a regulatory mechanism of NT5C2. A) Cartoon representation of the full-length NT5C2 in the inactive state (PDB code: 6DDQ). Subunits A and B are shown as cyan and yellow. The side chains of the amino acids forming the regulatory region (green) are depicted as stick models and labeled. The N-terminal region in subunit A is colored in pink and the side chains of important amino acids are shown with stick models. B. Cartoon representation of the full-length NT5C2 in the active state (PDB code: 6DE3). The view and the color codes are similar to A). The N-terminal helices, α1 and α2, are labeled. C) Immunoblot analysis of Jurkat cells expressing empty vector (EV) or FLAG-tagged mutants of NT5C2. D) Viability assay of Jurkat cells infected with empty vector or FLAG-tagged mutants of 470-478 region of NT5C2 lentiviruses treated with vehicle or increasing doses of 6-MP. Graphs show mean ± SD of three independent experiments performed in triplicate. E) Viability assay of Jurkat cells infected with empty vector or FLAG-tagged mutants of 470-478 region of NT5C2 lentiviruses treated with vehicle or increasing doses of 6-MP. Graphs show mean ± SD of three independent experiments performed in triplicate. *P* values were calculated using IC50 values and a two-tailed Student t test over wild type.

## Discussion

New genomic variants associated with cancer are being continuously identify. However, the functional contribution of these variants to oncogenesis, tumor progression, clinical outcome or response to therapy remain unknown. Notably, the computational and experimental methods for investigation of gain-of-function are limited. Altogether, our work demonstrates that structural methods can improve the identification and characterization of gain-of-function mutations in cancer. In particular, we have found that these mutations tend to occur in the interface between ordered and disordered protein regions, especially those that contain a functional site for protein-protein interaction or post-translational modification. Besides applications on oncogenes, our method can be applied to all genes to identify their functional regulatory regions. Compared to previous cancer mutation hotspot prediction methods, our method does not use the tumor mutation recurrence data as features or training labels, thereby excluding the possibility of information leakage when developing or evaluating our method. Comparing to the spectrum of novel unsupervised methods based on multiple sequence alignment or protein language models, the addition of structure-based information improves the interpretability of our score. We make available the codebase and prediction results of all human genes on (https://github.com/fuxialexander/ES).

Our study also highlights the importance of protein-protein complex structures to characterize the functional impact of protein mutation. As demonstrated in our analysis of NT5C2, ES scores that based on the predicted dimer structure leads to better concordance with the saturated mutagenesis experiment than monomer structure. This indicates that functional structure and conformation is crucial for studying mutation impact on proteins. Comprehensive structure prediction efforts targeting known pairwise protein-protein interactions have already begun^28,29^, which will provide rich data for our methods in the future. We believe these structures will provide clearer definitions of protein function and mutational mode of action, adding another milestone in the road from genotype to phenotype.

A corollary of these findings Is the Identification of new regulatory regions of NT5C2. NT5C2 relapse-associated gain-of-function mutations alter intrinsic switch-off mechanisms, blocking the enzyme in the active configuration and conferring resistance to 6-MP chemotherapy^26^. Importantly, the identification of new regulatory domains will provide novel insight on the NT5C2 enzymatic control and in mechanisms of 6-MP resistance in ALL. Interestingly, no mutations have been identified in these regions in ALL patients. However, it has been reported that nongenetic mechanisms such us post-translational modifications, affect NT5C2 activation and become drivers of therapy resistance^30^. In addition, our results demonstrate the effect of altering these regions in the response to 6-MP. Therefore, we cannot rule out the potential implication of these and other domains in the regulation of the NT5C2 enzymatic activity and as drivers of 6-MP chemotherapy resistance in relapsed ALL.

## Materials and Methods

### Data acquisition

#### Cancer somatic mutation data

We downloaded the complete gene somatic mutation data table from COSMIC (February 2015 version)^21^, where each row describes a single mutation in a specific patient. We kept only variants annotated as missense variants on protein product of canonical transcripts of all genes. In total, the collected mutation data covers 2,833,545 residues on 19,431 genes. We define the residues with more than 10% of all variant recurrence in each protein as its mutation hotspots.

#### Alphafold predicted structures

We downloaded apo-structure prediction of human reference proteome from the Alphafold database^20,31^. pLDDT where extract from the PDB files based on the b-factor value annotation. Cα-distance between each residue is computed for all structure using biopython^32^, resulting in a pairwise distance matrix D for each protein.

To focus on interacting residues only, we applied a user defined distance cutoff (20 °A used throughout this paper) to the matrix.

#### Short-linear motifs and protein-protein interaction interface annotation

We downloaded list of known short-linear motif (SliM) region from PhosphoSitePlus^33^. The SliM list contains 14,460 regions in 5,639 proteins from UniProtKB^34^. We also obtained human protein-protein interaction (PPI) interface from Interactome INSIDER^17^, which contains 14,445 proteins and 110,207 pairwise PPI with known interface residues in at least one of the interacting proteins. Overlapping with our oncogene COSMIC mutation list, we annotated 66,157 mutated residues (from which we count 7,882 in ‘Interface’, 2,786 in ‘SliM’ and 1,061 in ‘SliM & Interface’) in 270 oncogenes.

#### Oncogene annotation

We acquired a list of oncogene from the OncoKB^35^ Cancer Gene List (http://oncokb.org/cancerGenes) by keeping all genes with ‘Oncogene’ or ‘Oncogene/TSG’ annotation. The gene list was then intersected with the COSMIC mutation data and Alphafold database, resulting in 292 genes for downstream analysis.

#### Mutation probability analysis

To evaluate the probability of mutation from a specific codon (e.g. CGA) to a particular amino acid caused by cancer-specific mutagenesis process, we collect all COSMIC somatic mutations and calculate the probability of mutation happen to a particular codon. As a visual reference, we mark the top 10% residues with highest mutation probability using a grey dot in the ES-p plot like Figure 1A. A full visualization of mutation probability of NT5C2 is shown in Supplementary Figure 2.

### ES score construction

#### Evolutionary component

We compose the evolutionary component from a protein language model called Evolutionary Scale Modeling (ESM)^19^. The ESM model is trained on 98 million protein sequence across the evolutionary history to capture the (co)evolution constraint among residues. Taking a protein sequence input, the model is able to project the sequence into a high dimensional space that can indicate functional properties and structural dependence features of the protein. Specifically, we use the ‘Wildtype marginal probability’ score generated by a zero-shot version of the model, ESM-1v^13^. For wild-type residue *I* mutated to amino acid *mt* from wild type amino acid *wt*, the score is defined as:

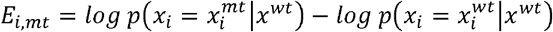

where *x_i_* is the marginal probability of a certain residue *i* given the sequence context around *i*.

We compute the evolutionary component using all 5 trained ESM-1v model and use the average output to reduce technical noise. Note that due to current technological and computing limitation, the ESM-1v model is hard to scale up to proteins with more than 1,024 amino acids. We bypass this limitation by breaking proteins longer than 1,000aa into 500aa segments and getting the score separately for each segment.

#### Structural component

The structure component is extracted from the residue-level predicted confidence from AlphaFold, namely the pLDDT score. As mentioned in the main text, we use the first derivative of the pLDDT score to detect change in prediction confidence. Specifically, we first smooth the pLDDT score using a Gaussian kernel (width=10aa) along each protein sequence, then compute the squared first derivative of the smoothed score. To remove the outlier caused by disordered region on both ends of the protein sequence, we clip the squared derivative score of each protein to its 20-80 percentile to produce the final structural component of ES score. Dimer structure of full length wildtype NT5C2 was predicted using Colabfold^36^ with pretrained AlphaFold-multimer^37^ weights and default parameters.

#### Position-specific ES scores

For visualization purpose, we define a position-specific ES score (ES-p) by first computing the average evolutionary component across all 20 possible amino acids, then multiplying it with the structure component. The resulting scores was then subject to smoothing across physically interacting residues and min-max scaling to form the ES-p score. Specifically, the ES-p score is defined as:

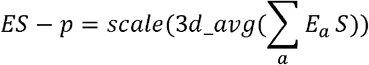

where *a* represents 20 possible amino acids and the function *3d_avg* is defined with an interacting distance threshold *D* < *d* is defined as

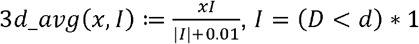

#### Mutation-specific ES scores

To produce a ES score not only specific to position but also to specific amino acid changes, we define mutation specific ES score (ES-mut) by adapting the practice established in a previous pan-cancer driver mutation study^8^. With a set of mutation data (specific cancer or pan-cancer), we first collect the ESM1v and ES-p score of all mutations, perform a principal component analysis and then use the scaled first principal component as the ES-mut score.

### Benchmark

To benchmark our method with previous methods on predicting oncogenic mutations, we downloaded the TCGA benchmark dataset from a pan-caner and pan-software study^8^ and selected mutations in oncogenes. To benchmark our method against EVE^12^, a new unsupervised variant pathogenicity prediction model, we downloaded all EVE prediction results from its website (https://evemodel.org/download/bulk), and identified overlapped genes and mutations with the benchmark dataset above. In total, our final benchmark dataset contains 5956 mutations in 107 oncogenes.

### Analysis of acute lymphoblastic lymphoma somatic mutation data

Somatic mutation and identified mutation hotspots in diagnostic and relapsed pediatric and adult acute lymphoblastic lymphoma (ALL) samples were acquired from a previous longitudinal ALL study^23^. Mutations on non-canonical protein isoforms that is not presented in AlphaFold database or UNIPROT are removed, affecting the following proteins: ABCA8, ABL1, ASXL2, AUTS2, CENPE, CHD4, CREBBP, DIAPH1, DNM2, DPP10, DSP, EDA2R, ERBB3, ETV6, FBN3, FBXW7, FHOD3, HTR3A, IGFN1, IKZF1, IL17RA, IL32, MYC, MYO9A, NIN, NIPBL, NOS1, NR3C1, PIK3C2G, PLCE1, PTPN13, PTPRB, PTPRS, RUNX1T1, SAMD4A, SETD2, TBX15, TENM2, TP53, TRPM3, UNC13A, VWF, WT1, ZAN, ZNF534. When computing the recurrence, we consider both clonal-wise and sample-wise mutations, that is, we identify unique mutations by DNA position, mutation allele, and sample ID.

#### Cell culture

We cultured Jurkat T cell line (American Type Culture Collection, ATCC) in RPMI-1640 containing 10% FBS, 100 U ml^-1^ penicillin G, and 100 100 μg ml^-1^ streptomycin. We cultured the HEK293T cell line (ATCC) in DMEM supplemented with 10% FBS, 100 U ml^-1^ penicillin G, and 100 μg ml^-1^ streptomycin. We performed cell culture in a humidified atmosphere at 37°C under 5% CO2 and we regularly tested for mycoplasma contamination.

#### Plasmid and vectors

We obtained pCDH-CMV-MCS-EF1α-Puro lentivector from System Biosciences (#CD510B-1). To generate the pCDH-NT5C2 expression construct, we PCR amplified NT5C2 fragment from pLOC-NT5C2 plasmid^38^ using the following primers: pCDH-NT5C2_Fw TTTGACCTCCATAGAAGATTATGTCAACCTCCTGGAGTG and pCDH-NT5C2_Rv AGCGATCGCAGATCCTTCGCTTATTCTTCCTCCTCCTCC. Fragments were then gel-purified and subjected into Gibson assembly (NEBuilder HiFi DNA Assembly Master Mix, New England Biolabs, #E2621) together with XbaI-NotI digested pCDH backbone. The product was used to transform NEB5a/C2987 competent cells. pCDH-NT5C2 plasmid was extracted from ampicillin-resistant colonies and confirmed by Sanger sequencing. We generated NT5C2 hydrophobic pocket (P472Q, F473P, R478H) and N-terminus mutants (Y27M, R34K, V37W) from pLOC-NT5C2 plasmid by site directed mutagenesis using the QuikChange II XL Site-Directed Mutagenesis kit (Agilent Technologies) according to manufacturer’s guidelines.

#### Lentiviral production and infection

We transfected lentiviral plasmids together with gag-pol (pCMV Δ R8.91) and V-SVG (pMD.G VSVG) expressing vectors into HEK293T cells using PEI transfection reagent (Polysciences, #24765). We collected viral supernatants after 48 h and used them to infect the Jurkat human cell line by spinoculation with 4 μg ml^-1^ Polybrene Infection/Transfection Reagent (Fisher Scientific). We selected infected human cell lines with 1 mg ml^-1^ blasticidin (InvivoGen, #ant-bl-1) for 14 days or with 1 mg ml^-1^ puromycin (Sigma-Aldrich, #P8833) for 7 days.

#### Drugs

We purchased 6-mercaptopurine monohydrate (6-MP, #AC226520050) from Thermo Fisher.

#### *In vitro* cell viability and chemotherapy response assays

We analyzed chemotherapy responses of human leukemia cell lines expressing wild-type NT5C2 or NT5C2 mutant constructs following 72-hour incubation with increasing concentrations of 6-mercaptopurine by measuring metabolic reduction of the tetrazolium salt MTT using Cell Proliferation Kit I (Sigma-Aldrich, #11465007001) following manufacturer’s instructions.

#### Saturation mutagenesis library

NT5C2 saturation mutagenesis synthetic library was created with an on-chip silicon-based DNA writing platform (Twist Bioscience), as previously described^39^. Briefly, oligonucleotide primers with mutations coding for all NT5C2 possible amino acidic substitutions and including, were synthesized on a silicon chip in equimolar ratios. Primers were then extracted, mixed and assembled to enable rapid production of high-diversity, full-length gene fragments by overlap extension PCR assembly. We next assembled full-length NT5C2 gene fragments with flanking sequences that contained overlapping regions with XbaI-NotI digested pCDH-CMV-MCS-EF1α-Puro vector, using Gibson assembly (NEBuilder HiFi DNA Assembly Master Mix, New England Biolabs, #E2621). Gibson reactions were pooled and transformed into E. coli XL-Gold cells. Final pCDH-NT5C2 saturation mutagenesis plasmid library was extracted by maxiprep.

#### Saturation mutagenesis screen

We infected Jurkat cells with pCDH NT5C2 saturation mutagenesis library at a multiplicity of infection of 1 and selected infected cells by treating with 1 μg ml^-1^ puromycin for 7 days. We removed dead cells via Ficoll density gradient centrifugation. We next treat triplicate cultures containing 500x representation of the library with vehicle-only or 6-mercaptopurine at the calculated 7-d IC90. We again removed dead cells via Ficoll density gradient centrifugation and extracted genomic DNA with phenol-chloroform using standard procedures. We next amplified full-length 1.7kb NT5C2 DNA insert by PCR using KAPA HiFi HotStart ReadyMix PCR Kit (Kapa Biosystems, #KK2601) and the following primers: *NT5C2sm_Fw* GTGATGCGGTTTTGGCAGTACA and *NT5C2sm_Rv* GTGATGCGGTTTTGGCAGTACA. We purified PCR product using 0.5x AMPure XP (Thermo Fisher Scientific, #NC9933872). Then, we sonicated DNA amplicons into ~300bp fragments using a S220 focused-ultrasonicator (Covaris) and we generated indexed libraries for next-generation sequencing using iDeal Library Preparation Kit (Diagenode, #C05010020), following manufacturer’s guidelines. Then, we sequenced these on an Illumina MiSeq instrument, using MiSeq Reagent Kit v2 (500-cycles) (Illumina, #MS-102-2003) for pairend sequencing DNA libraries loaded with a 20% spike-in of PhiX DNA.

#### Saturation mutagenesis analysis

The sequencing reads were mapped to NT5C2 cDNA sequence. For each residue, all codon changes included in the library design were identified in the mapped reads and counted. To distinguish a true amino acid change from sequencing errors, background noise was modeled by aggregating negative binomial distributions^40^. Briefly, if the total number of reads for a locus in sample *I* is *N_i_* among which *n_i_* reads harbor the amino acid change of interest, the distribution of *n_i_* follows a binomial distribution given *N_i_* and *a priori* probability of occurrence as error. When *N_i_* is sufficiently large (i.e., *n_i_*<<*N_i_*), the posterior predictive *p* value for having detected a true mutation in a test sample given the data observed in control samples can be approximated by the negative binomial distribution, followed by multiple hypotheses correction and false discovery rate (FDR) analysis^41^. Using this approach, FDR was calculated for each residue change in individual replicates from treatment experiments by comparison against the wild-type samples. FDR < 1e^-4^ was considered significant. Number and density substitution panels were generated in R using glot and ggplot2 packages.

#### Western blot analysis

We lysed cells in RIPA buffer, cleared of cell debris and boiled with 1x SDS-loading buffer. We performed BCA protein quantification according to manufacturer guidelines (BCA Protein Assay Kit, Fisher Scientific) and we loaded equal amounts of lysate onto a 4-12% Bis-Tris gel (Life technologies), separated by SDS PAGE, and transferred to a nitrocellulose membrane for western blot analysis. We detected NT5C2 with mouse anti-NT5C2 (Sigma Aldrich, #WH0022978M2) and rabbit anti-pSer502-NT5C2 (dilution 1:1000) (Covance) antibodies and β-actin with a mouse monoclonal anti-β-actin antibody (Sigma Aldrich, #A5441). We visualized the immunoblots using an Odyssey Infrared Imaging System (LI-COR Biosciences). We performed quantification analyses of the immunoblots using Image Studio software (LI-COR Biosciences).

#### NT5C2 structure analysis

Crystal structure representations were generated using PyMOL (http://www.pymol.org). We use fulllength NT5C2 double mutant (R39Q, D52N) in the inactive state (PDB code: 6DDQ) and in the active state (PDB code: 6DE3).

#### Statistics and reproducibility

We conducted statistical analyses using Prism software v8.0 (GraphPad software) and considered statistical significance at P < 0.05. We reported results as mean ± s.d. with significance annotated by P value calculated as indicated in the figure legends using Student’s t-tests assuming equal variance and normal distribution. The investigators were not blinded to allocation during the experiments and outcome assessment. The experiments were not randomized. No data were excluded from the analyses.

## Author Contributions

A.F. and R.R. designed the study. X.F. and R.R. designed the algorithm. C.R., R.A., J. W. L., H. K. and A.F. carried out the experimental systematic characterization of NT5C2 mutants. V. S. critically read the manuscript. All authors contributed to the manuscript.

## Acknowledgements

We gratefully acknowledge funding from NIH (R35CA253126 to RR, P01 CA174653 to RR, R01 HL159377-to A.F. and R.R., ADD) and SU2C Convergence 3.14 to RR. CR is supported by a Leukemia & Lymphoma Society Special Fellow award. RA is supported by the Leukemia & Lymphoma Society Postdoctoral Fellowship award.

---

**Supplementary Figure 1.**
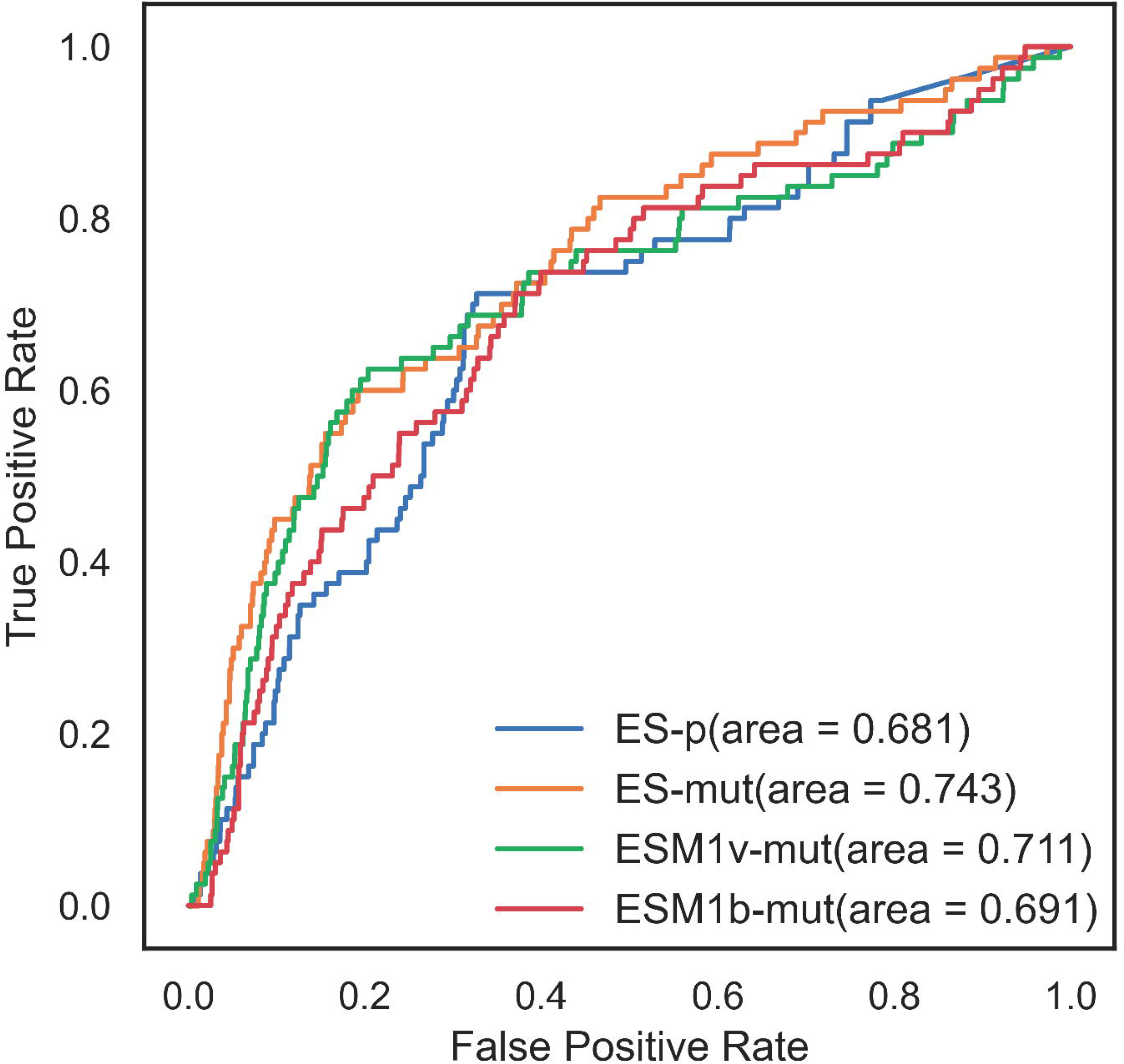
Comparison of different unsupervised methods for predicting hotspot mutation in oncogene. ROC curve for position-specific ES Score (ES-p), mutation-specific ES Score (ES-mut), mutation-specific scores from ESM1v (ESM1v-mut) and ESM1b (ESM1b-mut) model. Area under curve is marked in the legend. Higher area under curve means better prediction performance.

**Supplementary Figure 2.**
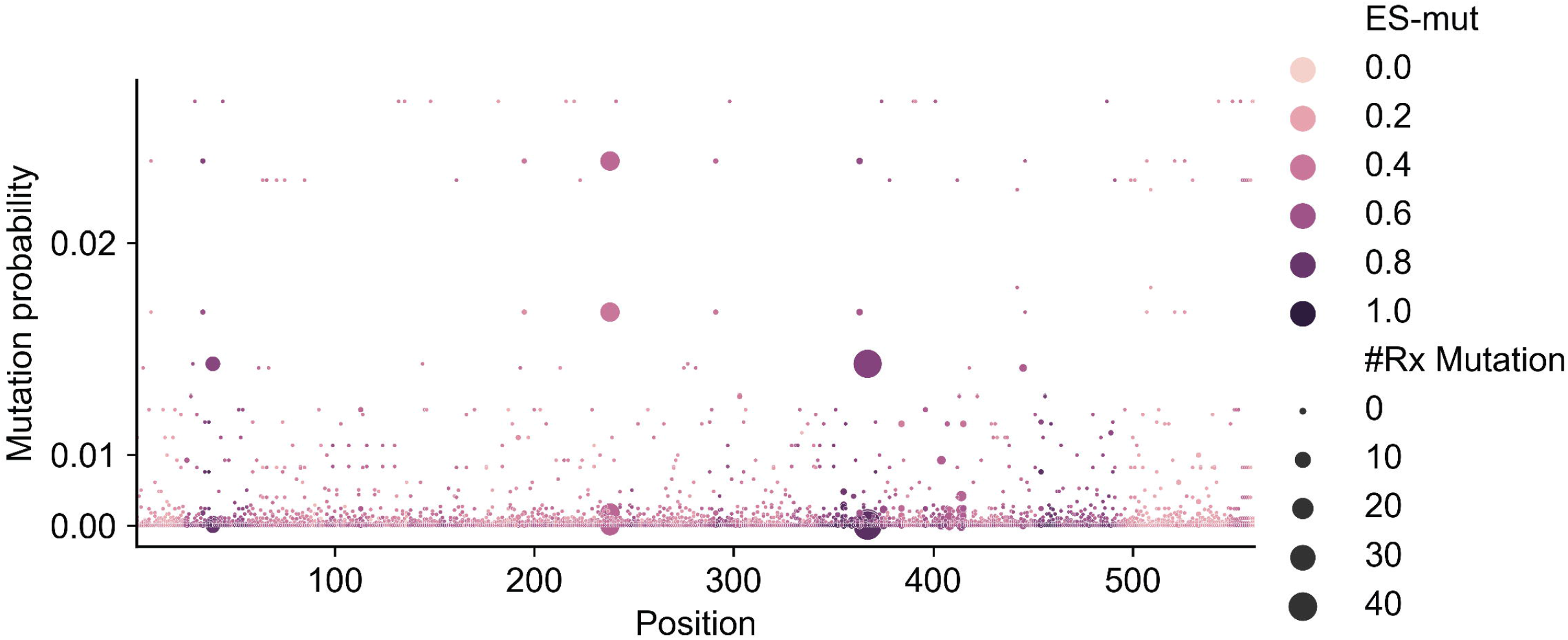
Mutation probability profile of NT5C2. Mutation probability (y-axis), ES-mut score (color) and mutation count in relapsed ALL samples (point size) were ploted along the protein sequence (x-axis). The 450-500aa region have high ES score but only limited number of high mutation probability points, corresponding to it’s paucity in mutation count.

**Supplementary Figure 3.**
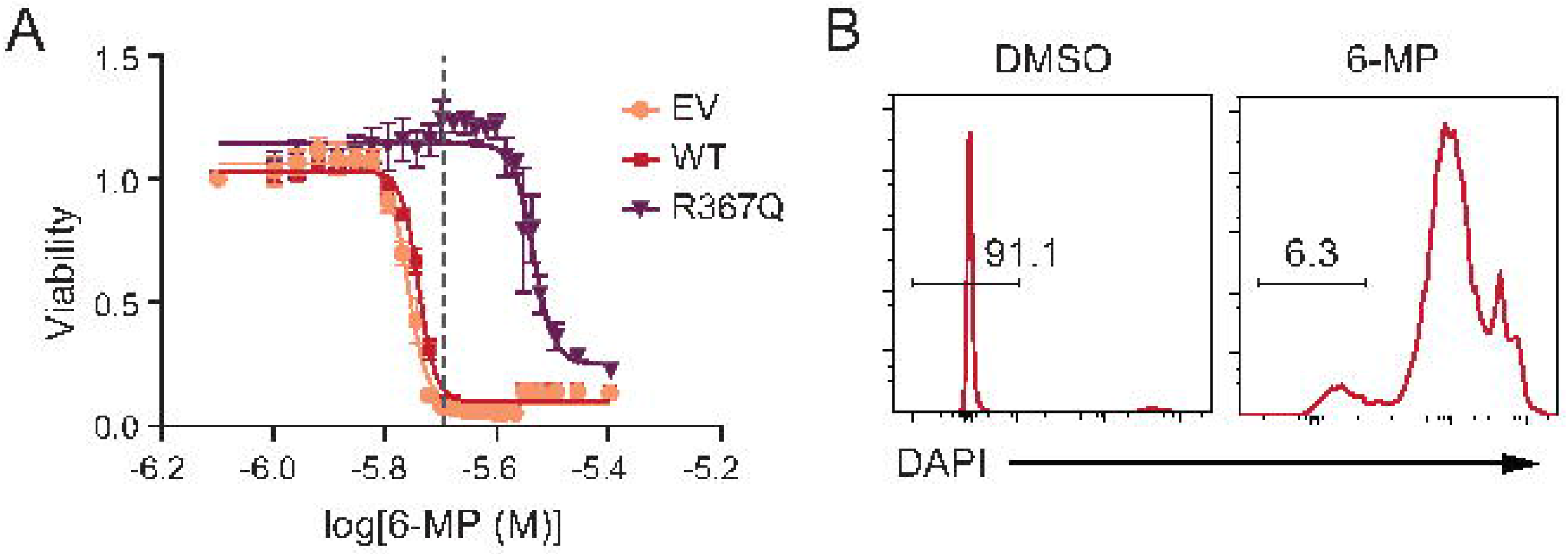
Selection of 6-MP resistant variants in the NT5C2 saturation mutagenesis screening. A) 7-day viability assay of Jurkat cells treated with vehicle or increasing doses of 6-MP. Graphs show mean ± SD of three independent experiments performed in triplicate. B) Flow cytometry viability analysis of Jurkat cells after at the calculated 7-d IC90 6-MP treatment using DAPI as marker. Percentages of live cells are shown.

